# Exact calculation of the joint allele frequency spectrum for isolation with migration models

**DOI:** 10.1101/065003

**Authors:** Andrew D. Kern, Jody Hey

**Author notes:** Corresponding Author: Andrew D. Kern Department of Genetics Rutgers University 604 Alison Rd. Piscataway, NJ 08854.

## Abstract

Population genomic datasets collected over the past decade have spurred interest in developing methods that can utilize massive numbers of loci for inference of demographic and selective histories of populations. The allele frequency spectrum (AFS) provides a convenient statistic for such analysis and accordingly much attention has been paid to predicting theoretical expectations of the AFS under a number of different models. However, to date, exact solutions for the joint AFS of two or more populations under models of migration and divergence have not been found. Here we present a novel Markov chain representation of the coalescent on the state space of the joint AFS that allows for rapid, exact calculation of the joint AFS under isolation with migration (IM) models. In turn, we show how our Markov chain method, in the context of composite likelihood estimation, can be used for accurate inference of parameters of the IM model using SNP data. Lastly, we apply our method to recent whole genome datasets from African *Drosophila melanogaster*.

## INTRODUCTION

The explosion in availability of genome sequence data brings with it the promise that longstanding questions in evolutionary biology might now be answered. In particular, understanding the balance of evolutionary forces when populations begin to diverge from one another is crucial to our understanding of the process of speciation. Population genomic sampling of multiple individuals from closely related populations provides our clearest view of the evolutionary forces at work during divergence, however it remains a challenge as to how best to analyze such massive datasets in a population genetic framework (Sousa and Hey 2013).

A popular model for population divergence is the so-called isolation with migration (IM) model (Wakeley 1996; Nielsen and Wakeley 2001; Hey and Machado 2003), in which a single ancestral population splits into two daughter populations at a given time and the daughter populations then have some degree of geneflow between them. IM models are a convenient framework for statistical estimation of population genetic parameters as the models described by various parameter combinations exist along a continuum between pure isolation after divergence to panmixia among daughter populations. More complex models of divergence, for instance secondary contact after isolation or geneflow that stops after a certain period of time, are also readily modeled in the IM framework. As a result numerous methods are now available for estimation of IM parameters.

Generally there exist two classes of methodology for the estimation of IM model parameters: genealogical samplers which aim to accurately compute the probability of a population sample under the assumption of no recombination within a given locus (e.g. IMa2; Hey and Nielsen 2007; Hey and Nielsen 2004) and methods which make use of the joint allele frequency spectrum (AFS) and assume free recombination between SNPs (e.g. δaδi; Gutenkunst et al. 2009). While genealogical samplers yield maximum likelihood or Bayesian estimates of population parameters, they become somewhat unwieldy for use with genome-scale data, due to the assumption of no recombination. Thus with the enormous increase in population genomic data from both model and non-model systems, much recent effort has been devoted to AFS based approaches that rely upon composite likelihood estimation (Gutenkunst et al. 2009; Naduvilezhath et al. 2011; Lukić and Hey 2012; Excoffier et al. 2013).

Estimation methods based on the joint AFS between populations center around calculating the probability of an observed AFS given the vector of parameters that describe the population history. The method for calculation of this expected AFS is thus central, and varies between competing methods. For instance, Gutenkunst et al. (2009) took the approach of numerically solving a diffusion approximation to the population allele frequency spectrum, whereas more recent methods of demographic inference rely upon coalescent simulation to estimate the expected sampled AFS (Naduvilezhath et al. 2011; Excoffier et al. 2013). While both of these approaches have been shown to be reliable for demographic inferences under many parameterizations, both are approximate and may contain error to various degrees across parameter space.

Here, we introduce a method for exact calculation of the joint AFS under two-population IM models with continuous migration. Our method uses a coalescent Markov chain approach that is defined on the state space of the AFS itself. Using this newly defined state space, in combination with the rich mathematical toolbox of Markov chains, we are able to readily compute the expected AFS of a given IM model for moderate sample sizes (say *n*_1_ = *n*_2_ *<* 9). We compare our coalescent Markov chain calculations of the AFS to diffusion approximations and that obtained via simulation. Further, using simulation we show how our approach can be used for accurate inference of demographic parameters. Lastly we apply our software package implementing the method, IM CLAM, to population genomic data from African populations of *Drosophila melanogaster*.

## MODEL

Here we present a strategy for exact calculation of the joint AFS under the IM model, and the subsequent inference of its associated parameters, that relies upon both discrete time and continuous time Markov chains (DTMC and CTMC respectively). In outline our approach involves first enumerating the complete state space associated with a given configuration of samples from two populations (i.e. sample sizes), followed by construction of a transition matrix to be used for a DTMC (or the analogous CTMC), and finally through the use of standard Markov chain techniques, the calculation of the implied joint AFS. For reasons that will become clear below, we begin by describing how one would calculate the exact joint AFS from a two population island model, before moving on to the full-blown IM model.

### A markov chain on the state space of the joint AFS

The first step in our approach requires the complete enumeration of the state space associated with our Markov chains given a sample configuration. The state space we describe is on the space of the allele frequency spectrum. That is to say that each state of our model implies a unique contribution to the joint AFS of the model in question. To track the allele frequency contribution implied by each state we will track the number of descendent leaf lineages in each population that each gene copy present is ancestral to. We will need to track this quantity independently for each population to deal with migration. To introduce our state space consider a sample that consists of one allele for population 1 and one allele from population 2, and let *n*_1_ and *n*_2_ be the sample sizes such that *n*_1_ = *n*_2_ = 1 (Figure 1). Although this is a trivially small case, it is adequate for accurately describing the form of the state space. Our initial state (i.e. the configuration at the time of sampling), call it *A*_0_, is

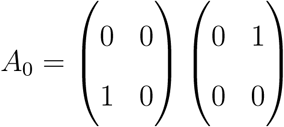

**Figure 1:**
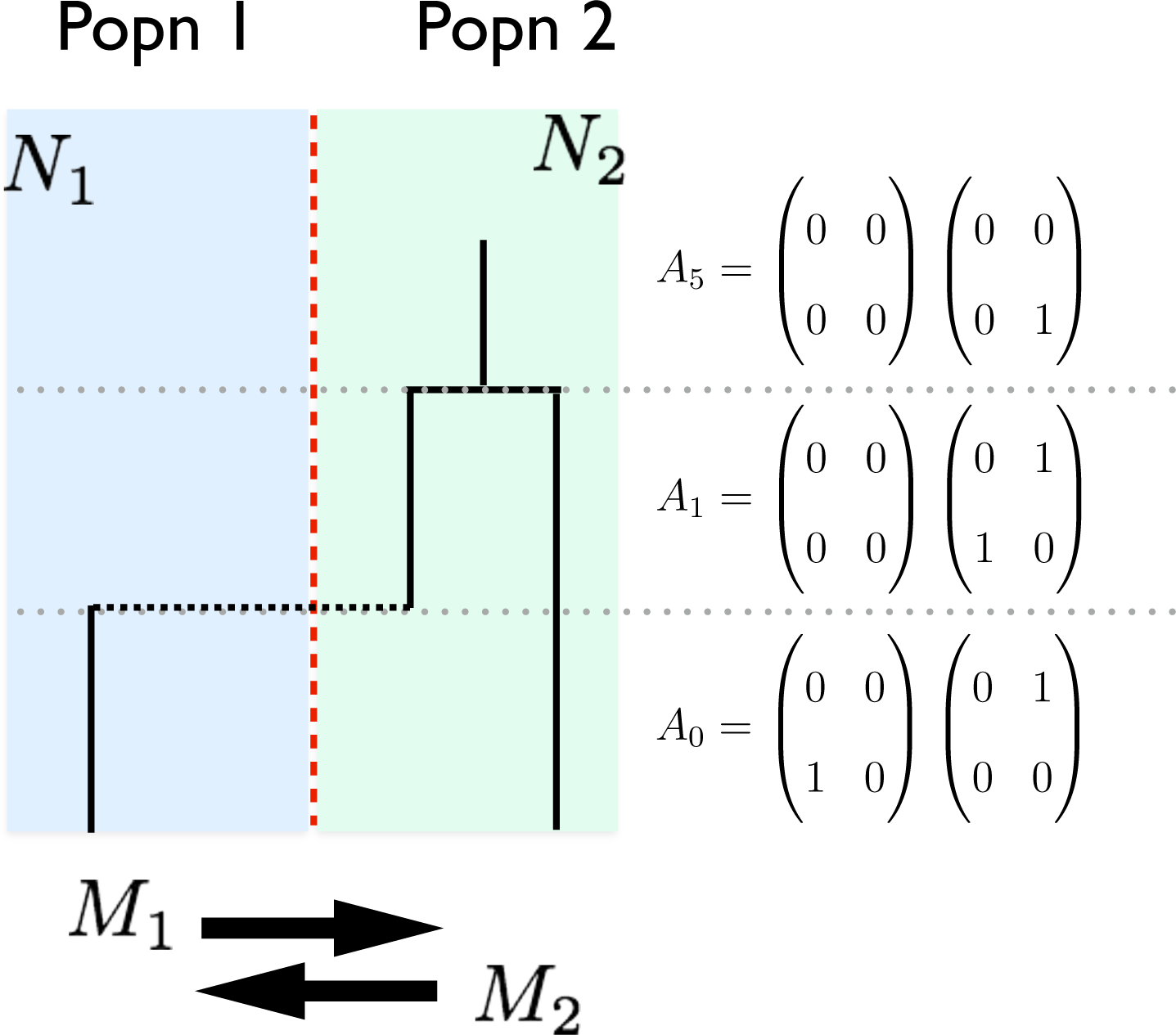
Representative State Space for a two population Island Model. hown here is an example of a two population island model with sample sizes *n*_1_ = 1 and *n*_2_ = 1. The model has two population sizes, *N*_1_ and *N*_2_, and two migration rates, *m*_1_ and *m*_2_. The representative state space at each phase in the coalescent tree is shown to the right. 31

where the left and right matrices represent the state in populations 1 and 2 respectively, and the entry at *i, j* represents the number of gene copies ancestral to *i* sampled alleles in population 1 and *j* sampled alleles in population 2. By convention these state space matrices are zero indexed, and there will never be a non-zero value at the position (0,0) as the model does not track lineages that are not ancestral to the sample. The initial state *A*_0_indicates that there is a single allele in population 1 that is ancestral to one of the sampled gene copies from population 1 and a single allele in population 2 that is ancestral to one of the sampled gene copies from population 2. Moving back in time in Figure 1 the first event is a migration event from population 1 to population 2. Thus in state *A*_1_ the matrix representing population 2 now has two alleles, one of which is ancestral to the sampled gene copy in population 1 and another that is ancestral to the sampled gene copy from population 2. Further notice that the left hand matrix, representing population 1, is empty. Finally to two alleles coalesce to find the MRCA in population 2, as indicated in state *A*_5_.

To enumerate the complete state space associated with a given sample configuration (*n*_1_, *n*_2_), we use a recursive approach that considers all possible coalescent and migration moves among present gene copies to exhaustively find all possible states, including MRCA states that will represent the absorbing states of our Markov chain. Note that in this two population island model only two absorbing states are possible–the MRCA could be found in population 1 or it could be found in population 2. In the case of *n*_1_ = *n*_2_ = 1 as shown in Figure 1, there are a total of 6 possible states however the number of states grows extremely quickly with increasing sample size (See Appendix). For instance when *n*_1_ = *n*_2_ = 2 there are 46 possible states, and *n*_1_ = *n*_2_ = 3 there are 268 states. Figure 2 shows how the state space grows in sample size, and while growth is sub-exponential it clearly explodes for larger samples. In Figure S1 we show the associated compute time to calculate the AFS of an IM model as a function of sample size using the implementation introduced below. Computational complexity grows quickly in sample size, indeed nearly exponentially, so we suggest limiting use of our method to samples of size *n*_1_ = *n*_2_*<* 9, although larger samples would be feasible on appropriate hardware.

**Figure 2:**
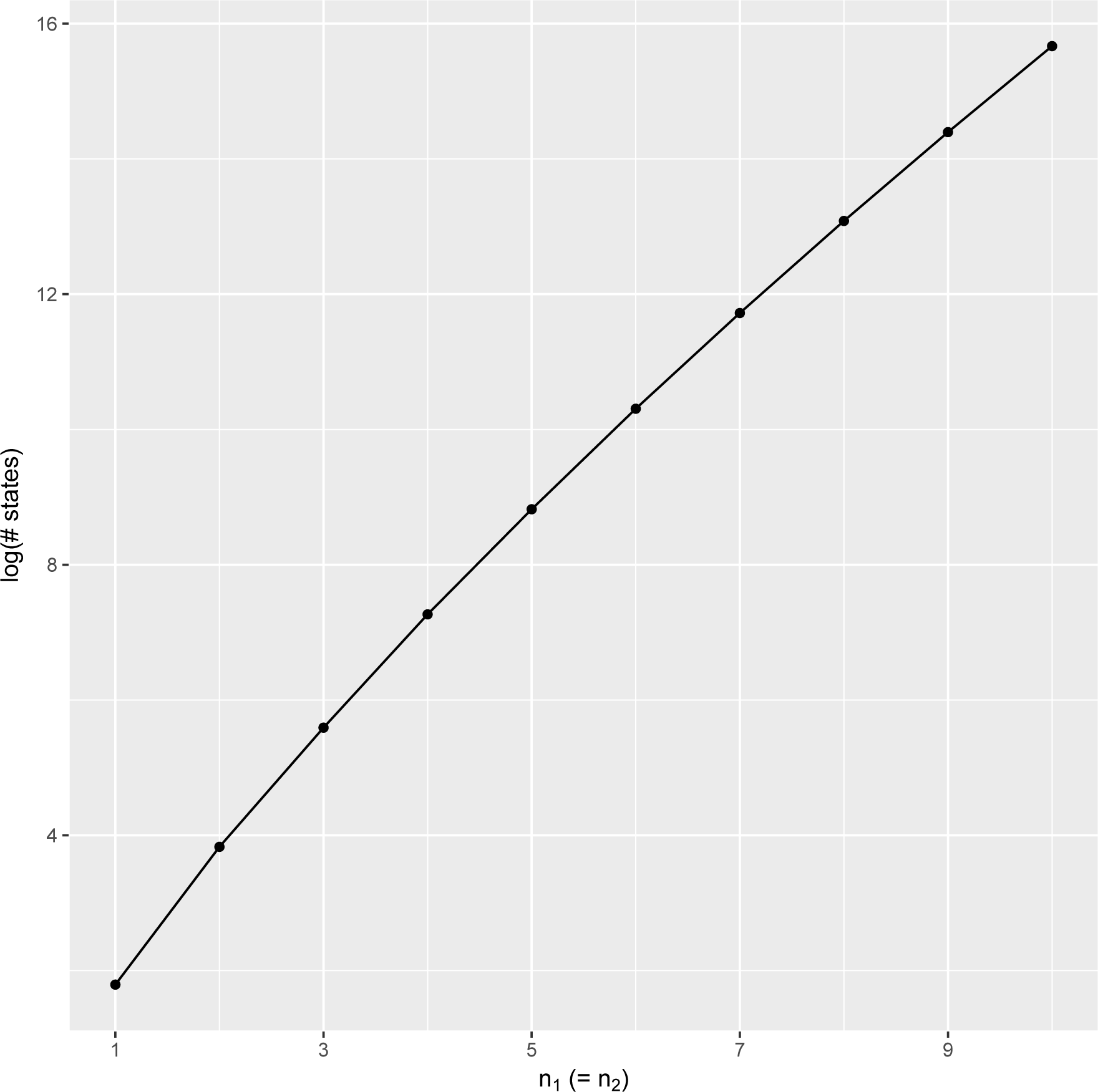
State space expansion as function of sample size. Here we show how the state space grows as function of sample size considering symmetric sampling such that *n*_1_ = *n*_2_. Note that the y-axis is shown on a log scale.

### Markov chain transition matrix

Having defined the state space we next consider the form of the transition matrix associated with the DTMC. Transitions between states in our coalescent markov chain depend both on parameters of the model (e.g. population sizes, migration rates) and on the combinatoric probability involved in the chain move. For instance let *n_i_* be the number of active lineages in population *i* within a state of the chain. Further let *N_i_* be the population size of population *i* and its associated coalescent rate be *C_i_* = (*n_i_* − 1)*/*4*N_i_*. Finally define *M_i_* as the migration rate from population *i* scaled by effective population size such that *M_i_* = 4*N_i_m* where *m* is the fraction of the focal population made up of migrant individuals each generation. Jumps of the chain will depend on these parameter dependent rates (*C_i_*, *M_i_*) as well as the combinatoric probabilities which only depend on the configuration of lineages present within each state.

Consider first the probability of a coalescent event in population *i* that moves the chain, *{ζ}*, from state *A_y_* at time *t* to *A_z_* at time *t* + 1. Such a coalescent event could happen either between two lineages of different types (i.e. ancestral to different numbers of sampled gene copies among populations) or between two lineages of the same type (i.e. ancestral to the same numbers of sampled gene copies among populations), so let us label our two focal lineages that will coalesce as *k* and *l*. If *k* and *l* are of the same type and if there exist *x* copies of this lineage type in population *i* within state *A_y_*, then the combinatoric probability of such an event, call it *T*(*A_y_*, *A_z_*), would be 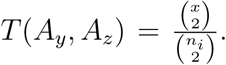 If *k* and *l* are of different types and there exist *x_k_* copies of *k* and *x_l_* copies of *l* in population *i* within state *A_y_*, then 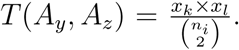 We can now write down the complete probability of a coalescent event as

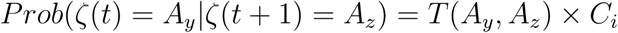

Notably the first combinatoric term, because it does not depend on parameters of the model, can be pre-calculated and the transition matrix simply updated by scaling by the coalescent rates of interest.

In the case of a migration event from population *i* to *j* the terms of the transition matrix take the form

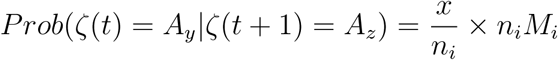

where as before *x* is the number of copies of the lineage involved in the migration defined by the transition from *A_y_* to *A_z_*.

To make this concrete, lets focus for a moment on the state space of a sample of size *n*_1_ = 2, *n*_2_ = 1. This complete state space is included in the appendix and labeled B. Next consider the coalescent event that transitions the chain from state 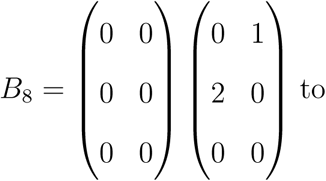 to state 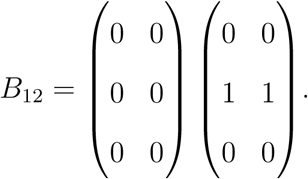 In this move one of the two lineages that are only ancestral to gene copies from population 1 coalesces with the lineage that is only ancestral to population 2. Using the equation above the combinatoric probability of the move is 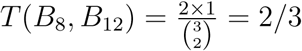 and the complete probability of the transition between states is *Prob* (ς(*t*) = *B*_8_*|ς*(*t* + 1) = *B*_12_) = *T*(*B*_8_, *B*_12_)*× C*_2_ = 2*/*3 *×* 3*/*2*N*_2_ = 1*/N*_2_. If we consider instead the other possible coalescent event from state *B*_8_ that moves the chain to state 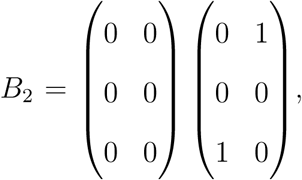 then two lineages of the same type have coalesced, 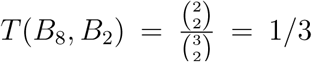 and *Prob*(ς(*t*) = *B*_8_*|ς*(*t* + 1) = *B*_2_) = *T*(*B*_8_, *B*_2_)*× C*_2_ = 1*/*3 *×* 3*/*2 *N*_2_ = 1*/*2 *N*_2_.

Turning our attention back to the case of *n*_1_ = *n*_2_ = 1 (see appendix), the unnormalized transition matrix associated with the DTMC, call it *P*, would be

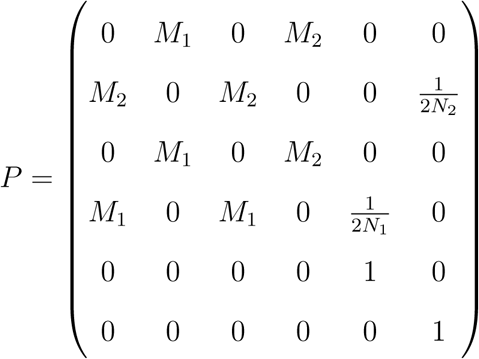

where now each matrix entry *P_ij_* is scaled so that each row sums to one such that Σ*_j_ P_ij_* = 1. Each *P_ij_* represents the probability of the Markov chain moving from state *A_i_* to state *A_j_* in the next jump. The *P* matrix also implies an analogous CTMC transition matrix, call it *Z*, whose rows are constrained such that Σ*_j_ Z_ij_* = 0. With these transition matrices in hand we now turn attention to computing the SFS of the island (or IM) model.

#### Calculating the AFS

As said above, each state implies an associated contribution to the allele frequency spectrum. Let *F* represent the joint AFS from a two population sample. *F* will be matrix valued of size *n*_1_ + 1 rows and *n*_2_ + 1 columns, where *n*_1_ and *n*_2_ are the sample sizes from populations 1 and 2 respectively. Entries of *F*, *F_ij_*, will be the number of SNPs sampled with *i* derived alleles in population 1 and *j* derived alleles in population 2. To map a given state *A_i_* to its contribution to *F*, we need only ask how long the system stays in a given state (i.e. the expected duration) and then add that amount of time to each of the corresponding cells of *F* from the non-zero entries in both the right and left hand matrices of the *A_i_th* state. This is justified as the probability mass associated with each cell of the AFS is simply proportional to the mean total length of branches that when mutated lead to frequencies of the focal AFS position when normalized by the mean total length of the associated coalescent tree (Adams and Hudson 2004).

We can use the tools of Markov chains to then perform the two calculations needed to exactly calculate the AFS under a given model: 1) calculate the expected number of times each state is visited before absorption (i.e. reaching the MRCA), and 2) calculate the expected length of time the chain is in each state to compute the AFS. The latter calculation is simply the exponentially distributed waiting time under the coalescent with migration, which itself is a function of the number of gene copies active in a given state, population sizes, and migrations rates.

Calculating the expected number of visits to each state is more involved. We can rearrange our transition matrix *P* into what is called “canonical form”. We assume that *P* has *r* absorbing states and *t* transient states, such that

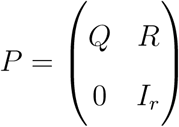

where *Q* is a *t × t* submatrix, *R* is a *t × r* submatrix, and *I_r_* is the identity matrix of rank *r* (Kemeny and Snell 1976). Using this factorization we can next compute the fundamental matrix of our Markov chain, *N*, by using the relationship

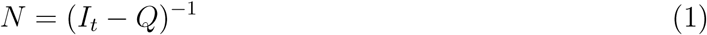

where the entries *N_ij_* represent the expected number of visits to state *j* given the chain started at state *i*, and *I_t_* is a rank *t* identity matrix. It is important to note that this calculation will thus require the inversion of a potentially very large matrix, thus complicating our implementation. For the calculation of the island model however, we are only interested in one row of *N*, as the starting state is known with certainty (i.e. the observed sample), so this is readily solved. Also, note that *N* gives us the expected number of visits to each state by the DTMC until absorption (i.e. the MRCA). For the island model this describes the complete stochastic process as in that case we are dealing with a time homogenous process. For models with changes in population size or populations splitting we would have to consider different “phases” of the demographic history separately, as the transition rates through the system, or indeed even the state space of the system will change moving back in time.

Returning for a moment to the island model then, having calculated *N* we are ready to compute the expected AFS. As we said before, the expected AFS will simply be the sum of the products of the number of visits to each state and the length of time spent in each state. For the island model in the case where *n*_1_ = *n*_2_ = 1 there will be 6 terms in the summation to find *F*, one for each state.

#### Isolation with Migration

To calculate the AFS for the IM model, we calculate the contributions to the AFS from two sources: that of the island model phase of the model prior to divergence (looking back in time), and the contribution to the AFS from the single, ancestral population (see Figure 3). The contribution to the AFS from the island model portion, call it *F_I_*, can be computed by first calculating the total AFS from the island model from time zero to absorption, *F_tot_*, and then subtracting off the portion of the AFS contributed from the population divergence time, *t_div_*, until absorption (e.g. Wakeley and Hey 1997). Let the vector *π*(*t*) be the probability of being in each state of our Markov chain at time *t*. We need to calculate *π*(*t_div_* both to find *F_I_* and to figure out where our system begins the single population phase of the IM model. We use a CTMC representation of our same transition matrix from the island model (denoted *Z*) to compute *π* (*t_div_*) using the matrix exponential such that

**Figure 3:**
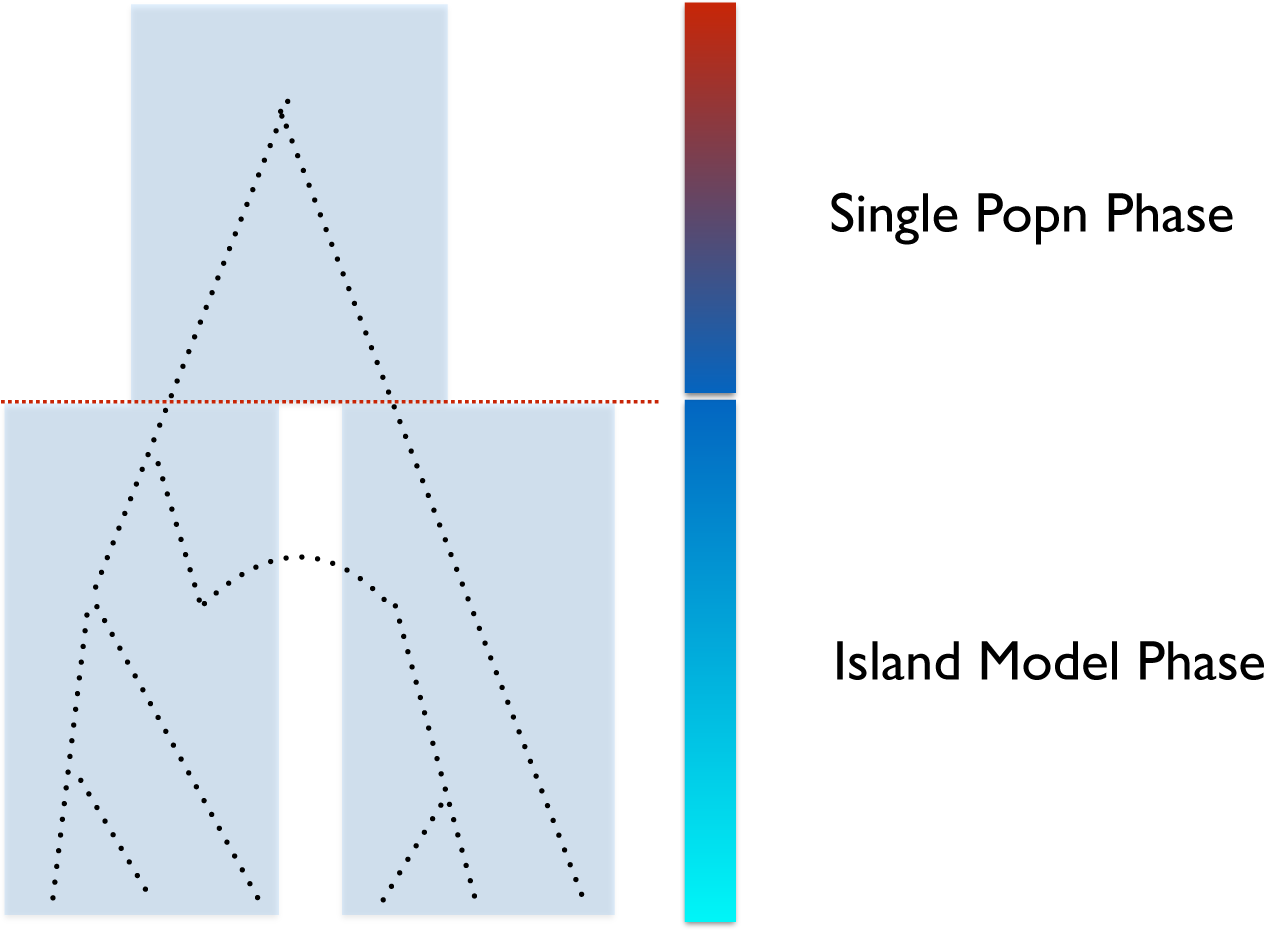
Two phases of the IM Model. Here we illustrate the two phases of the IM model, the first of which is an island model phase and the second, the ancestral single population phase. To compute the expected AFS of the IM model we calculate the AFS contributions from each of these phases separately and then combine them to get our AFS from the IM model.

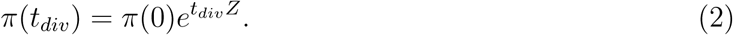

With *π* (*t_div_*) in hand, we can use the fundamental matrix of the island model, *N*, to compute the number of visits to each state conditional on starting in each state at *t_div_* with probability *π* (*t_div_*) as *N_g_* = *π* (*t_div_*) *N*, where *N_g_* is subscripted *g* in reference to the fact that these represent “ghost visits,” unseen in the actually IM model. *F_I_* then can simply be calculated as *F_I_* = *F_tot_* − *F_g_*, where *F_g_* is the AFS implied by *N_g_*.

Once we have the contribution to the AFS from the island phase, *F_I_*, there is only one portion remaining–the contribution to the AFS from the single population, ancestral phase, call it *F_A_* (Figure 3). To compute this we map the state space of the island model onto a reduced state space of a single population model, use that mapping to fold *π* (*t_div_*) to the state space size, and then compute a new DTMC transition matrix for the single population phase, changing population size as necessary and removing migration. With the new transition matrix we can compute the fundamental matrix for the ancestral phase, *N_A_*, and from that its contribution to the AFS, *F_A_*. Finally the AFS for the complete IM model, *F_IM_*, is equal to the combined sums of the AFS contributions from the two phases such that *F_IM_* = *F_I_* + *F_A_*.

### Composite Likelihood

Our goal is to calculate the probability of an observed AFS given a set of IM parameters. To do this we use the now familiar composite likelihood approach, treating individual SNPs as independent observations (Adams and Hudson 2004; Gutenkunst et al. 2009). We model SNPs as being drawn from a multinomial distribution with probabilities drawn from the joint AFS. Let *x* be a set of SNP frequency observations from two populations, and be the full set of parameters from the IM model. Then the probability of observing our data given the parameters, i.e. the likelihood of Θ is

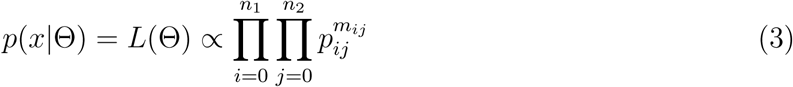

Here the indices *i* and *j* are over the domains of the AFS, and *m_ij_* is the observed number of SNPs in *i* individuals in population 1 and *j* individuals in population 2. The *p_ij_* terms are the entries of the exact expected AFS calculated as described above. Lastly, the likelihood above is proportional up to an appropriate multinomial constant which can be dropped from the likelihood calculation as it does not depend on the parameters.

## IMPLEMENTATION

Our strategy for computing the AFS from the IM model relies upon taking the inverse of two large, sparse matrices, corresponding to functions of the transition matrix from the DTMC, and exponentiating one matrix. Such calculations are extremely expensive computationally, so in our implementation of this method we have used parallel, scalable algorithms where ever possible. Our software package, IM CLAM (<u>I</u>solation with <u>M</u>igration via <u>C</u>omposite <u>L</u>ikelihood <u>A</u>nalysis using <u>M</u>arkov chains), performs these calculations with help from two open source packages, the CSPARSE library (Davis 2006) and the PETSc package (Balay et al. 1997; Balay et al. 2015a; Balay et al. 2015b). In particular we use PETSc to distribute all sparse matrix calculations across a parallel compute environment that uses MPI. For matrix inversion, we compute row by row of the inverse matrix using a direct solver from CSPARSE and distribute those solves across cores. The matrix exponential is calculated using the Krylov subspace method as implemented in the SLEPc add-on to the PETSc package (Hernandez et al. 2005). IM CLAM and its associated open source code are available for download from GitHub (https://github.com/kern-lab/im_clam).

Estimation of uncertainty surrounding our point estimates is implemented in our software package by solving for the Godambe Information Matrix (Godambe 1960). This is done by numerically calculating the inverse of the Hessian matrix for the likelihood function at the joint composite likelihood estimates of the parameters along with a variability matrix that examines the variance in the gradient of the likelihood across bootstrapped samples. Uncertainty estimation via the Godambe Information Matrix has recently been shown to be appropriate for composite likelihood estimation where it replaces the more familiar Fisher Information Matrix (Varin et al. 2011; Coffman et al. 2016).

## APPLICATION TO DROSOPHILA MELANOGASTER DATA

We apply our method to recent whole genome sequencing projects from *Drosophila melanogaster* in which multiple smaller population samples from a variety of African populations have been sequenced to good depth (Pool et al. 2012; Lack et al. 2015). We obtained aligned datasets from the Drosophila Genome Nexus resource (v1.0; Lack et al. 2015), and subsequently filtered from those alignments regions that showed strong identity-by-decent (IBD) and admixture using scripts provided with the alignments. From these we chose a subsample of 5 populations with sample sizes that were small enough for efficient estimation using IM CLAM. On our compute hardware, IM CLAM estimation for samples of size *n* > 9 was prohibitively slow, we thus chose the following populations to analyze: Dobola, Ethiopa (ED; *n* = 8), Kisoro, Uganda (UG; *n* = 6), Maidiguri, Nigeria (NG; *n* = 6), Donde, Guinea (GU; *n* = 7), and Phalaborwa, South African (SP; *n* = 7). The joint AFS was then constructed for each pairwise combination, using alignments to *D. simulans* and *D. yakuba* to determine the derived and ancestral allele at a given SNP. Tri-allelic positions were ignored. In an effort to sample the AFS from regions of the genome that should be less likely to affected by linked selection, we only examined intergenic regions that were at least 5 *kb* away from genes, and that did not contain simple repeats, repeat masked regions, annotated transcription factor binding sites, or annotated regulatory elements. This yielded 5530 regions of the genome with a total length of 4.43Mb. For each population pair we performed three parameter optimizations from different starting conditions and verified that all optimizations converged to the same estimates. For estimation of uncertainty we used the Godambe Information Matrix calculated using 100 bootstrap replicates from the observed AFS. Run times on 96 cores of Xenon 2.5gz processors varied between 4 and 14 hours.

## RESULTS

### Simulation

We first set out to compare the expected AFS calculated with IM CLAM versus that calculated from coalescent simulations. As our calculations result in the exact AFS, we were interested in comparing the convergence of the simulated AFS to the true AFS as a function of the number of simulations. In Figure 4 we show the mean percentage error of the AFS computed from simulating a given number of independent genealogies with a small mutation rate (*θ* = 0.001) and rejection sampling only those trees that contained SNPs. In Figure 4 the AFS was computed using *n*_1_ = *n*_2_ = 6, a symmetric migration of rate *m*_12_ = *m*_21_ = 1.0, and a divergence time of *t_div_* = 0.25. As the number of simulated genealogies increases the mean percentage error between the simulated AFS and that calculated by IM CLAM drops quickly. However after 10^6^ simulations the amount of Monte Carlo error plateaus at approximately 0.3% and then decays very slowly even after 10^9^ simulations. Thus brute force simulation of the AFS seems ill advised for IM models, as it will be computationally quite expensive to converge to the correct distribution of allele frequencies, although approximately correct calculation could be done with considerably fewer simulations.

**Figure 4:**
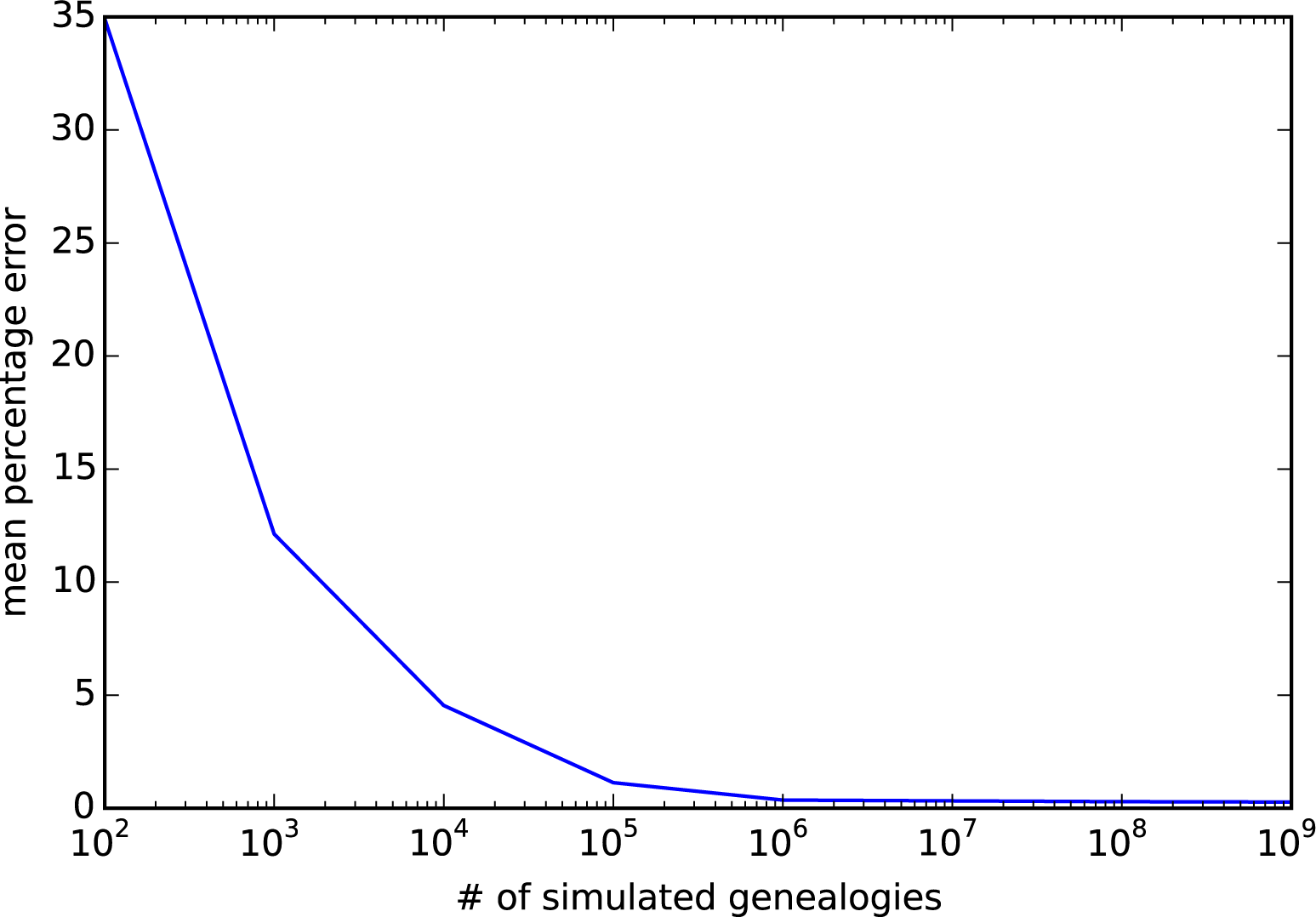
Monte Carlo error in simulations of the allele frequency spectrum. Here we show the decline in the mean percentage error in estimates of the joint AFS from simulations where we vary the number of independent coalescent genealogies simulated in comparison to our exact solution.

We next turned our attention to comparing our exact AFS to that computed by the popular software package *∂a∂i* (Gutenkunst et al. 2009). *∂a∂i* uses diffusion approximations to model the joint AFS among two populations and thus itself may be susceptible to a certain amount of error for given parameterizations. We compared our exact AFS to that generated from *∂a∂i* under a range of migration rates, *m* = *{*0, 1, 5, 10*}*, and having fixed population sizes to 1.0 (IM CLAM considers each populations size relative to the size of population 1, where as *∂a∂i* normalizes by the size of the ancestral population) and *t_div_* = 0.5. Figure 5

**Figure 5:**
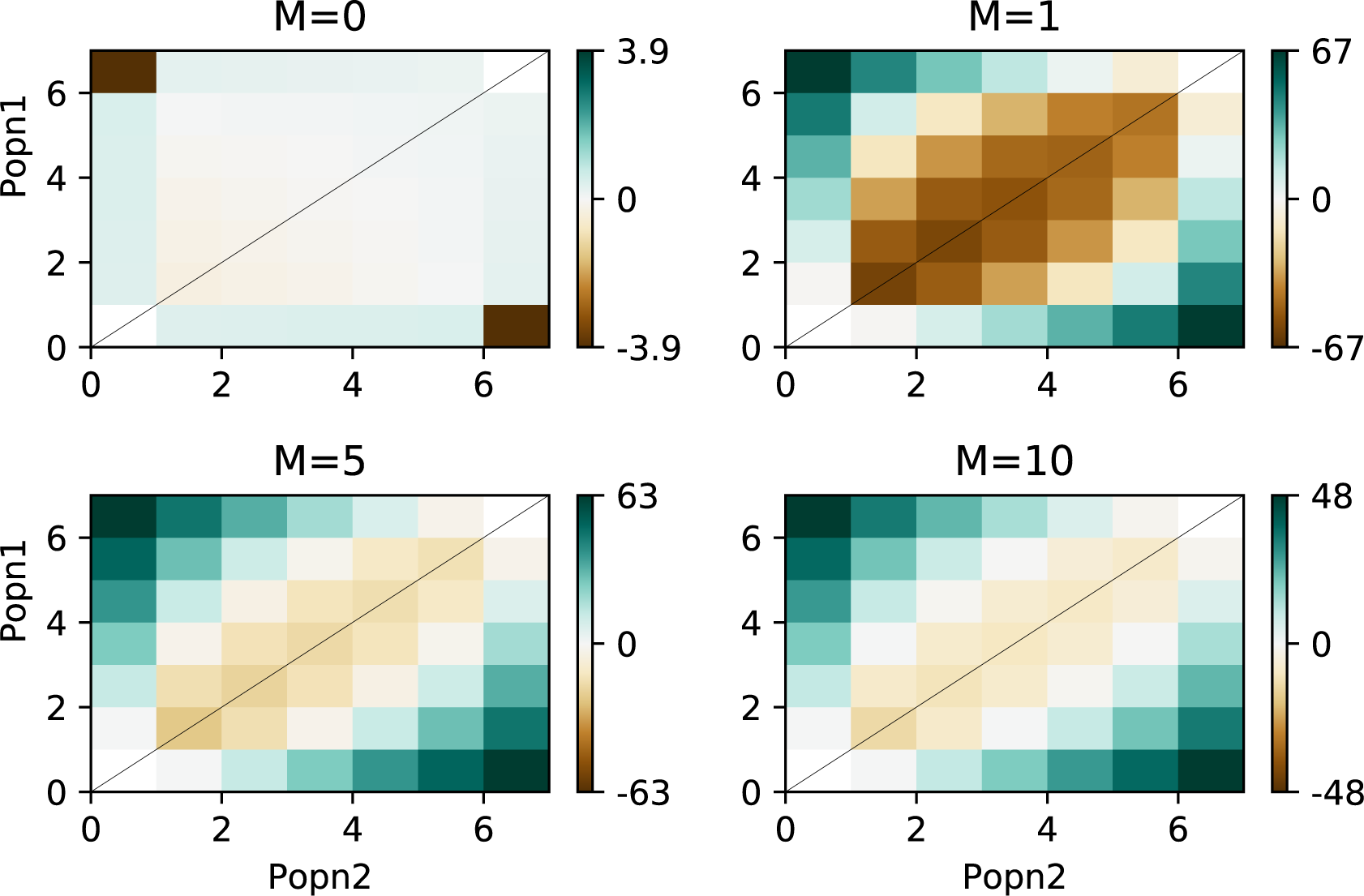
Percent deviation of expected AFS calculated from *∂a∂i*. Clockwise from top left panel we show the percent deviation of each cell in the expected AFS for four different symmetric migration rates *m* = *{* 0, 1, 5, 10*}* from *∂a∂i* versus our exact calculation.

shows the element wise percentage error for the *∂a∂i* approximation of this comparison. *∂a∂i* harbors an appreciable amount of error under these parameters, particular at the corners of the matrix, that represent fixed differences among populations. Thus while *∂a∂i* has been shown to be accurate for use in inference, we can see here that the expected AFS produced using the diffusion approximation still strays from the true value.

As a result of this discrepancy we set out to compare the accuracy of inference using IM CLAM in comparison to *∂a∂i*. Our goal here is not to perform an exhaustive comparison between methods, as IM CLAM is much more limited in scope than *∂a∂i*, however we wish to show that our method has utility for parameter inference as well. For this we generated 100 replicate simulated AFS draws using coalescent simulations in a manner as to simulate a large number of independent SNPs. Again we set *n*_1_ = *n*_2_ = 6, divergence time is one of *t_div_* = 0.1, 0.5, 1.0, and either a symmetric migration regime with rates *m*_12_ = *m*_21_ = 1.0 or asymmetric migration with rates *m*_12_ = 1.0, *m*_21_ = 0.1. We set a low per locus, *θ*, *θ* = 0.001, and generated 10^6^ genealogies. This yielded approximately 3.54 *×* 10^5^ SNPs per simulated AFS sample. With these simulated datasets we then set out to infer the parameters of the IM model. Figure 6 and Figure 7 show violin of parameter estimates for both IM CLAM and *∂a∂i* for symmetric and asymmetric migration respectively. Supplemental Figures S2 and S3 show root-mean-square error across all parameter estimates for the point estimates. In general both methods are relative accurate across parameterizations however it can be seen that a minority of optimizations using *∂a∂i* yielded outlier parameter estimates. From our simulated parameters, it seems that *∂a∂i* is most accurate at intermediate divergence times (*t_div_* = 0.5) and does less well under the other two divergence times simulated. In contrast IM CLAM performs well over all parameters considered here. It is worth considering that both methods are using the BFGS (Broyden-Fletcher-Goldfarb-Shanno; (Press 1985)) algorithm for optimization, set with the same stopping criterion and bounds on the parameter space explored, thus failed optimization alone seems an unlikely explanation. Indeed similar behavior for *∂a∂i* was observed in an earlier report (Naduvilezhath et al. 2011).

**Figure 6:**
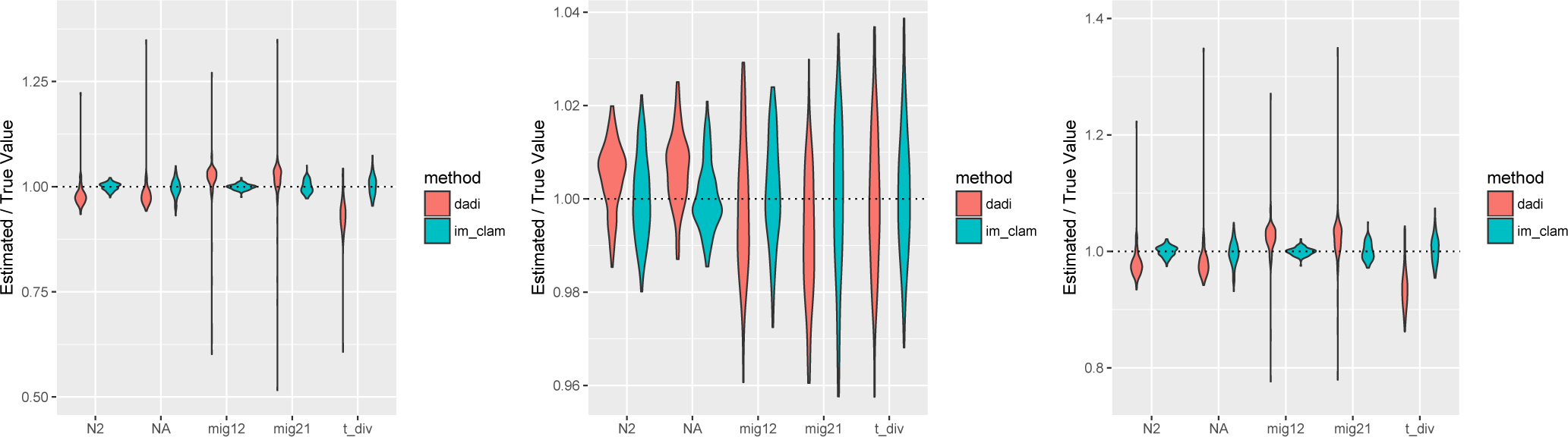
Accuracy of parameter inference using IM CLAM and *∂a∂i*. Shown are violin plots of point estimates from 100 replicate simulations with IM CLAM and dadi. Each panel uses the same migration rates, *m*_12_ = *m*_21_ = 1.0, but divergence time varies from left to right *t_div_* = 0.1, 0.5, 1.0.

**Figure 7:**
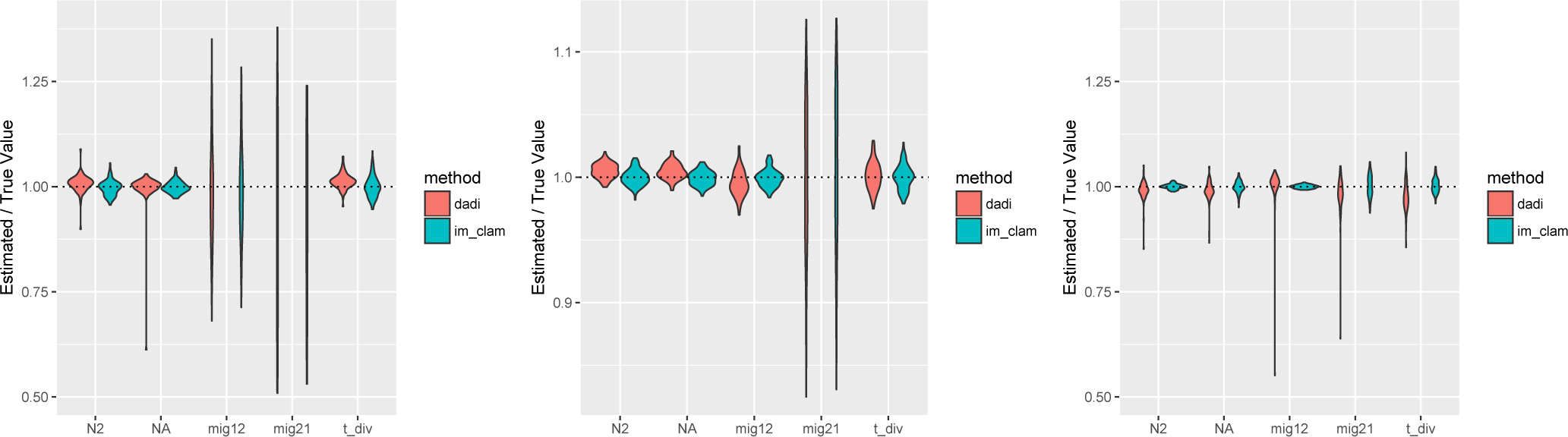
Accuracy of parameter inference using IM CLAM and *∂a∂i*. Shown are violin plots of point estimates from 100 replicate simulations with IM CLAM and *∂a∂i*. Each panel uses the same migration rates, *m*_12_ = 1.0; *m*_21_ = 0.1, but divergence time varies from left to right *t_div_* = 0.1, 0.5, 1.0.

### Application to *Drosophila melanogaster* data

The demographic history of *Drosophila melanogaster* in many ways mirrors that of human populations. *Drosophila melanogaster* is commonly thought to have had its origins in sub-Saharan Africa, and have spread out of Africa approximately between 10,000-20,000 years ago (Lachaise et al. 1988; David and Capy 1988; Begun and Aquadro 1993; Li and Stephan 2006). *D. melanogaster* seems to have first migrated to Europe and Asia via the middle east, presumably as a human commensal, and then only much later did it arrive in North America (Lachaise et al. 1988). While there is good genetic support for sub-Saharan Africa to be the ancestral range of the species (Begun and Aquadro 1993; Pool and Aquadro 2006) less is known about the history of populations within Africa. Levels of variation among populations do suggest that *Drosophila melanogaster* ancestrally occupied Southern Africa, and from there spread into western and northern Africa (Pool et al. 2012). Here we model the demography of five African populations, representing each of the major hypothesized axes of geographic range expansion. Using the joint AFS from each pair of these population samples we estimated IM model parameters using IM CLAM. Point estimates of pairwise population divergence time and its associated uncertainty are summarized in Table 1, and a UPGMA tree constructed from these population divergence times is shown in figure S4. These estimates are scaled in the number of individuals for population sizes and the number of years for divergence time by assuming a mutation rate per base per generation of *u* = 5.49 *×* 10^−9^ (Schrider et al. 2013) and 15 generations per year (Pool 2015). A complete table of parameter optimization results is given in Table S1. While the UPGMA tree is intended to be heuristic, care should be taken in its interpretation as a true, multi-population model has not been considered here, and instead we have reconstructed a tree based on pairwise divergence.

Population size estimates suggest that all sub-Saharan African populations have experienced significant population growth since their divergence from one another, with the exception of Ethiopia. Population growth varies between comparisons from 1.7x-3.8x depending on the specific population pair. This growth, because it is seen broadly across populations suggests a recent change in population size for the species, perhaps in the last few thousand years. The exception to this trend is Ethiopia, which appears to have undergone a significant bottleneck to between 0.57x-0.83x of its ancestral size depending on the pairwise comparison considered.

Estimates of divergence time point to the South African population representing an earlier lineage split among African melanogaster populations, with an average divergence between it and other populations of greater than 8,000 years (table 1; fig. S4). This is consistent with observations based on population differentiation that have suggested Southern African populations represent a possible ancestral population (Pool et al. 2012). Our tree based reconstruction of population divergence points to Nigeria being an outgroup to both Eastern African populations (Uganda and Ethiopia) as well as Guinea further to the west. The extent to which this can be interpreted to reflect the biogeographic history of the species in African is most likely limited due to the large levels of gene flow we estimate between most populations below.

**Table 1:** Divergence time estimates from *D. melanogaster* populations. Values below the diagonal are point estimates of *t_div_*, while above the diagonal 95% confidence intervals are given.

|  | SP | UG | ED | NG | GU |
| --- | --- | --- | --- | --- | --- |
| SP | - | $\pm 375.11$ | $\pm 254.84$ | $\pm 430.50$ | $\pm 358.13$ |
| UG | 9257.11 | - | $\pm 253.81$ | $\pm 525.89$ | $\pm 424.92$ |
| ED | 5577.46 | 3628.34 | - | $\pm 239.69$ | $\pm 255.89$ |
| NG | 10582.02 | 8325.86 | 4332.84 | - | $\pm 1122.92$ |
| GU | 9983.12 | 7203.86 | 3939.54 | 8496.81 | - |

Finally, our results point to broad, ongoing gene flow among African populations (Table S1). To visualize source-sink dynamics of gene flow among populations we present a circle plot of estimates of 4*Nm* in Figure 8. Figure 8 is scaled such that the width of each arc is proportional to 4*Nm* where *N* is that estimated from the focal sink population. A few general features can be gleaned from this plot. First, Ethiopia is the least well-connected population by migration per generation among the populations considered here. Second, South Africa is largely a sink population, rather than being the source of outgoing migrants. Third, Uganda, while it seems to be sending migrants to West and South Africa, receives fewer migrants proportionally than other populations. Lastly, Nigeria, Guinea, and Uganda seem to be potent sources of migrants both to one another as well as to South Africa.

**Figure 8:**
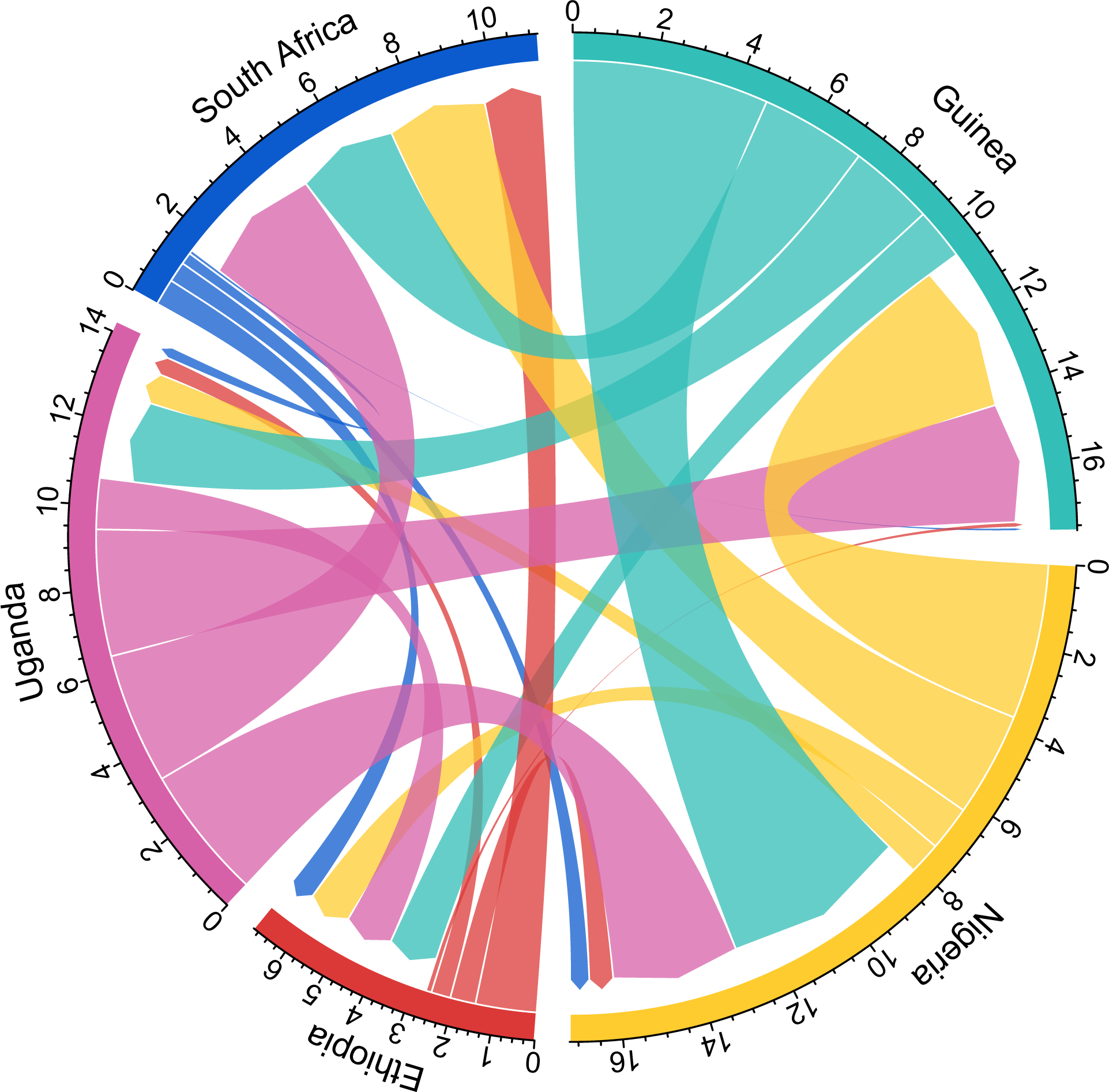
Relative migration rates between populations. Here we visualize relative rates of effective migration (4 *Nm*) between populations, show in units scaled by *N* of the sink population. Length of the outer arc associated with each population label gives a sense of the total flux into and out of each focal population.

## DISCUSSION

Population genetic inference of demographic history has become an increasingly important goal for modern genomics, as the impacts of demography on patterns of genetic variation is now appreciated to directly impair our ability to identify causative disease variation via linkage (e.g. Rogers 2014) as well as shape the genetic architecture of phenotypic variation within populations (Lohmueller 2014; Simons et al. 2014). Moreover, our understanding of human prehistory has been revolutionized in recent years through demographic inference using population genetic data (e.g Botigué et al. 2013; Ralph and Coop 2013; Raghavan et al. 2015; Poznik et al. 2016). While that is so, methods that efficiently utilize whole genome information for inferring rich demographic histories, particularly multiple population histories, still lag behind the huge availability of data (Sousa and Hey 2013). Accordingly, much recent effort has focused on using the joint allele frequency spectrum of samples drawn from multiple populations as a way to summarize genome-wide data for demographic inference (Gutenkunst et al. 2009; Naduvilezhath et al. 2011; Lukić et al. 2011; Lukić and Hey 2012; Excoffier et al. 2013; Kamm et al. 2015).

In this study we present a novel method for numerically calculating the exact joint allele frequency spectrum expected from two population Isolation with Migration models. Our method relies upon a Markov chain representation of the coalescent, in which the state space of the chain is the joint AFS at a given point in time. Through the use of this state space, in conjunction with standard Markov chain techniques, we are able to numerically calculate the exact expected AFS. Our method stands in contrast to other popular techniques that either use diffusion approximations (Gutenkunst et al. 2009; Lukić et al. 2011) or direct Monte carlo simulation (Excoffier et al. 2013) to estimate the expected AFS under a given parameterization. Indeed, as we have shown, estimation of the AFS via diffusion or Monte carlo simulation can lead to persistent error and in some cases numerical instability (see Kamm et al. 2015). While we here use a Markov chain approach to calculate the exact AFS under IM models, a recent, elegant paper by Kamm *et al.* (2015) presented analytic solutions and associated algorithms for computing the exact AFS for multiple population models with arbitrary population size histories but without continuous migration.

We have implemented our approach in a software package called IM CLAM that allows for inference of IM models using genome-wide joint AFS data by computing the exact AFS. As we have shown above with simulated data, IM CLAM is quite accurate in its inference of population parameters. Application of IM CLAM to population genomic data from *Drosophila melanogaster* populations sampled from sub-Saharan Africa points to a complex history of population divergence and ongoing gene flow among populations. Firstly, we find strong support for the notion that sub-Saharan populations, generally, have experienced population growth in the recent past and have not been at equilibrium for population size over an extended period of time. It is possible that such growth accompanied population expansion throughout the African continent from an ancestral range. Indeed our finding is consistent with earlier reports of population growth in African populations based on different population samples (Li and Stephan 2006; Sheehan and Song 2016). While this is so, our estimates of population growth occurring in the past few thousand years are much closer in line with what is reported for timing from Sheenhan and Song (2016) than from Li and Stephan (2006). The single exception to population growth, the Ethiopian sample, appears to have declined in size since its divergence from an ancestral population. Reduced population size of Ethiopia is corroborated by levels of heterozygosity observed in this population in comparison to other sub-Saharan samples (Pool et al. 2012).

Our results on population divergence times suggest that the South African population represents an ancient lineage that diverged from all other populations sampled greater than 8,000 years (fig. S4). This finding supports the hypothesis that Southern Africa might be the ancestral range of *Drosophila melanogaster*, in agreement with observations based on genetic differentiation and levels of heterozygosity (Pool et al. 2012). From this ancestral range it is likely that the species expanded first throughout west and central Africa, and only subsequently northward towards the horn of Africa. Decreased population size in Ethiopia is consistent with this scenario, suggesting a still-observable effect of a past population bottleneck in that sample. In general, the deeper divergence times estimated among African populations is striking–it seems that African populations have been diverging from one another for quite a long period of time. If we take at face value the biogeographic hypothesis that *Drosophila melanogaster* first expanded from Africa to Eurasia between 10,000-20,000 years (Lachaise et al. 1988; Stephan and Li 2007), divergence among African populations itself is only slightly less, mimicking to some extent the deep time population structure now believed to occur among some sub-Saharan human populations (e.g. Schlebusch et al. 2012).

While divergence time estimates are on the order of thousands of years among populations, estimates of gene flow suggest high ongoing rates of migration among many of the populations (fig. 8). Estimates of 4*Nm* among populations show considerable source sink asymmetry for the South African population, whereby the population appears to be taking in migrants but not sending them out. Ethiopia is also an outlier among sampled populations in our study for migration, as it appears to be the least well connected node in the network of geneflow through sub-Saharn Africa. Finally Nigeria, Guinea, and Uganda each are well connected via gene flow to each other and to lesser extents to South Africa. Inasmuch, while our estimates of population divergence suggest comparatively old split dates among populations, gene flow has been a potent homogenizing force among most of our sampled populations.

While the ability to compute the exact AFS under IM models using our Markov chain approach is an advance, there are many shortcomings to our methodology. Perhaps most challenging is the fact that the state space of our Markov chain grows nearly exponentially in sample size (fig. 2). This means that our approach is only computationally feasible for smaller sample sizes, as in the current state space the transition matrix associated with larger sample sizes will be too large to represent in memory, even when sparse matrix representations are used as we have done here. While this is so the state space of the Markov chain could potentially be reduced in size if by exploiting lumpability among states (*cf.* Andersen et al. 2014). Even at moderate sizes the computational costs of the matrix inversion and exponentiation needed by our method are still high, thus IM CLAM needs tens or hundreds of CPUs for optimization runs to complete within hours rather than days. To be applicable to larger samples then the user might proceed by taking subsamples of the AFS, for instance by projecting it down to the expected AFS given a smaller sample size using a hypergeometric distribution (Nielsen et al. 2005). Additionally our method could be extended to calculate the composite likelihood not just over sites, but over subsamples as well.

Despite the computational difficulties associated with the Markov chain approach described here, our method has opened a new avenue in calculating the likelihoods associated with AFS data and might be amenable to other population genetic problems. For instance, in the model presented above we consider the two dimensions of the state space matrices to represent different populations. It is simple to conceive of this dimension as instead two separate loci with recombination acting to make transitions among the numbers of alleles that are ancestral at one or both loci. In this way we have been able to write down a Markov chain that enables calculation of the two-locus allele frequency spectrum that itself might be useful for estimation of demographic parameters and recombination rates.

## ACKNOWLEDGMENTS

ADK and JH were supported by NIH R01GM078204. We thank Dan Schrider for suggesting the name IM CLAM, Yun Song for helpful conversations and comments about this effort, and two anonymous reviewers whose comments strengthened the paper.

## APPENDIX

The complete state space for a sample of configuration *n*_1_ = *n*_2_ = 1 is given below. The ordering of states shown is arbitrary but identical to the one used in the example markov chain transition matrix in the Model section of the paper.

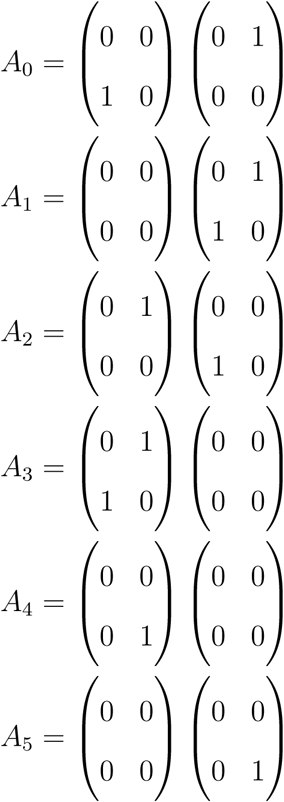

The next most simple state space is that for a sample of configuration *n*_1_ = 2, *n*_2_ = 1. This is referred to above in the Model section of the paper to illustrate the calculation of transition probabilities of the Markov chain. The complete state space in this case, call it B is given here

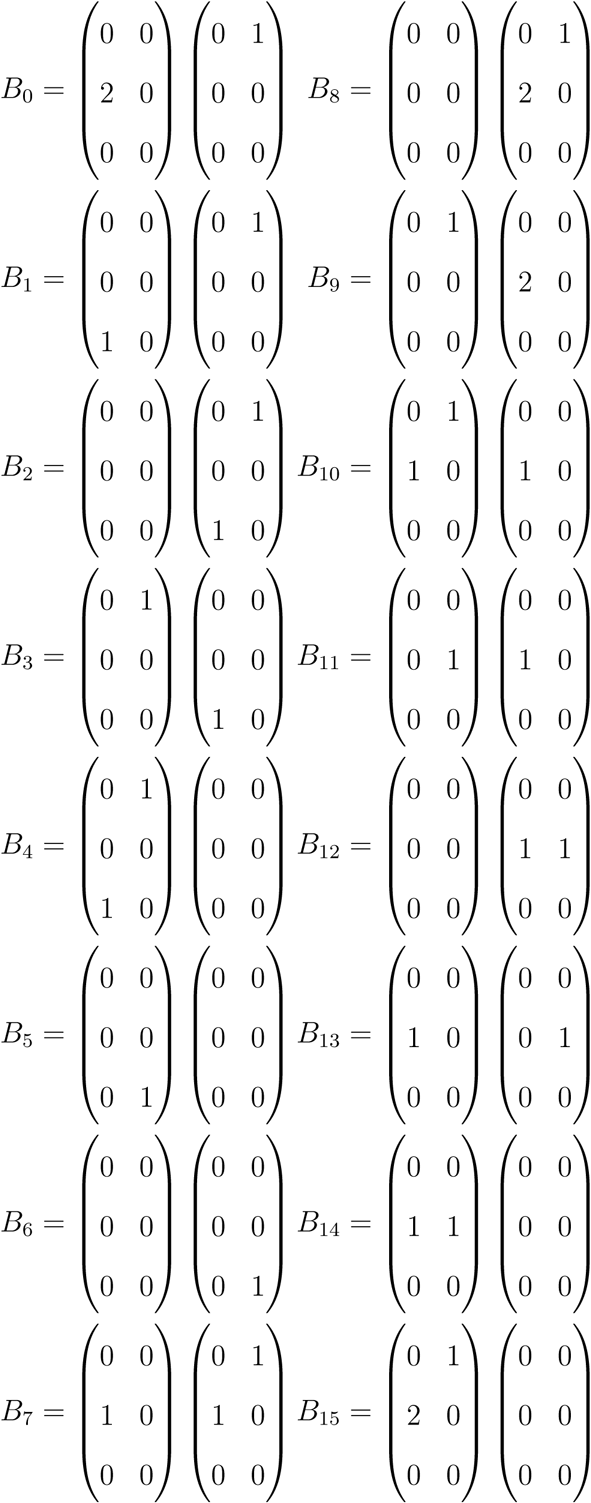

